# Intrinsic Neural Oscillations Predict Verbal Learning Performance and Encoding Strategy Use

**DOI:** 10.1101/2025.09.27.678958

**Authors:** Victor Oswald, Mathieu Landry, Hamza Abdelhedi, Sarah Lippé, Philippe Robaey, Karim Jerbi

## Abstract

Individuals adopt different encoding strategies to facilitate learning. However, few studies have investigated the neurophysiological basis that support these different encoding strategies across individuals. The present work addresses this gap by extending our previous findings on the direct relationship between cortical spectral power, measured via resting-state magnetoencephalography, and performance on standard cognitive test results. Our results highlight the complex interactions between endogenous brain oscillations, learning and verbal encoding strategies assessed by the California Verbal Learning Test (CVLT-2). First, we found that resting-state theta oscillations were significantly associated with verbal learning and subjective clustering strategies. Second, we observed that semantic clustering is facilitated by oscillatory patterns in left sensory-motor brain regions. Finally, our analyses revealed that serial and semantic clustering strategies are related to opposite regression patterns, indicating a competitive interaction. Together, these findings provide novel insights into the neural oscillatory dynamics that support diverse encoding strategies in verbal learning.

## Introduction

Spontaneous neuronal activity reflects the brain’s intrinsic functional organization and provides a powerful window into cognition ^1,2^. High-temporal-resolution techniques such as electrophysiology reveal that even in the absence of an explicit task, the brain exhibits complex, large-scale oscillatory patterns arising from neuronal synchronization ^3–6^. These patterns display specific frequency–spatial topologies shaped by neurophysiological pathways and evolutionary constraints and are linked to various cognitive abilities. For example, neurophysiological research has related fronto-parietal alpha power alpha peak amplitude at rest with global IQ, predicts working memory performance ^7,8^. Complementing these correlational findings, rhythmic TMS studies provide causal evidence by showing a frequency- and region-specific dissociation, whereby parietal alpha stimulation impaired working memory performance, whereas prefrontal theta stimulation supported prioritization processes ^9^. Such oscillatory signatures observed at rest represent a promising, though underexplored, avenue for investigating how intrinsic patterns of neural activity accounts for interindividual differences in cognitive performance. Resting-state activity may reveal neurophysiological predispositions that influence not only cognitive capacity but also the strategic approaches individuals naturally adopt when performing complex tasks. Prior studies have shown that spontaneous oscillations at the cortical level predict working memory performance and executive functioning ^7,10^. The present work aims to extend this line of research by investigating whether intrinsic brain organization at rest relates to both the efficiency of learning and the selection of specific encoding strategies in the context of memory formation.

Human memory capacity is constrained by the limits of short-term information storage ^11,12^. To overcome these limits, learners often rely on encoding strategies— cognitive operations that reorganize information to enhance retention ^13^. These include mnemonics, chunking, elaborate rehearsal, visual imagery, and semantic encoding. Such strategies may be self-initiated and idiosyncratic, reflecting stable interindividual differences ^14^. The neural correlates of semantic clustering, in particular, have been examined through neuroimaging, electrophysiology, and behavioral methods. Evidence shows that training participants to use semantic strategies increases activation in the occipital cortex, left orbitofrontal cortex, and left parietal cortex when recalling structured versus unstructured word lists ^15^. Explicit semantic instructions further enhance activation in the bilateral dorsolateral prefrontal cortex, inferior frontal gyrus, right orbital gyrus, occipital cortex, precuneus, and medial prefrontal cortex ^16–18^. In healthy adults, verbal learning and semantic clustering correlate with left hippocampal volume, while in individuals with depression, reduced hippocampal volumes—particularly on the right— are linked to greater reliance on serial clustering, a less efficient strategy ^19^. Deficits in semantic clustering following prefrontal lesions or in HIV-associated dementia underscore the importance of fronto-basal ganglia circuits in effective learning ^20–22^. While the role of the frontal cortex in semantic clustering is well established, the neural bases of other strategies remain less explored.

The present work investigates whether spontaneous oscillatory brain activity at rest can predict not only verbal learning performance but also the type of encoding strategy individuals naturally adopt. We used the California Verbal Learning Test (CVLT-2) ^23,24^, which measures verbal learning efficiency across five trials of word-list recall and quantifies three types of strategies: i) Semantic clustering: grouping items by category (e.g., fruits), ii) Serial clustering: recalling items in presentation order, and iii) Subjective clustering: recalling items in the order just produced.

Our aim was to identify RS oscillatory patterns that predict both performance and strategy selection, without explicit training or instruction, thereby capturing individuals’ innate tendencies. By mapping these resting-state predictors for grouping strategies, this study provides a neurophysiological framework for one of the most widely used clinical memory assessments, applied from childhood to aging ^25^.

## Methods

### Participants

Forty healthy subjects (n=40), comprising 25 females, with a mean age of 25.1 (± 4.64) years, participated in this study. None of them had a reported history of neurological or psychiatric disorders. The project received approval from the University of Montreal and the CHU Sainte-Justine Research Ethics Board. Informed consent was obtained from all participants before the experiment, and they were financially compensated upon completion.

### Behavioral Assessment

#### Neuropsychological evaluation

A comprehensive neuropsychological evaluation was conducted on the same day as the MEG recordings (Figure 1), employing the CVLT-2 ^26^ to assess verbal memory. This test includes two lists, each containing 16 words categorized into four semantic groups: transportation, vegetables, animals, and furniture. To minimize reliance on semantic or serial encoding strategies, consecutive words in each list were from different categories. Participants were instructed to recall the words in any order they preferred after each list presentation. Following the fifth presentation of the first list, a second (interfering) list was introduced. Subsequently, participants were asked to recall the initial list shortly after the second list presentation (short-term recall) and again following a delay of approximately 1 hour and 30 minutes (long-term recall). This delay was consistent for all participants and filled with other neuropsychological tests.

**Figure 1.**
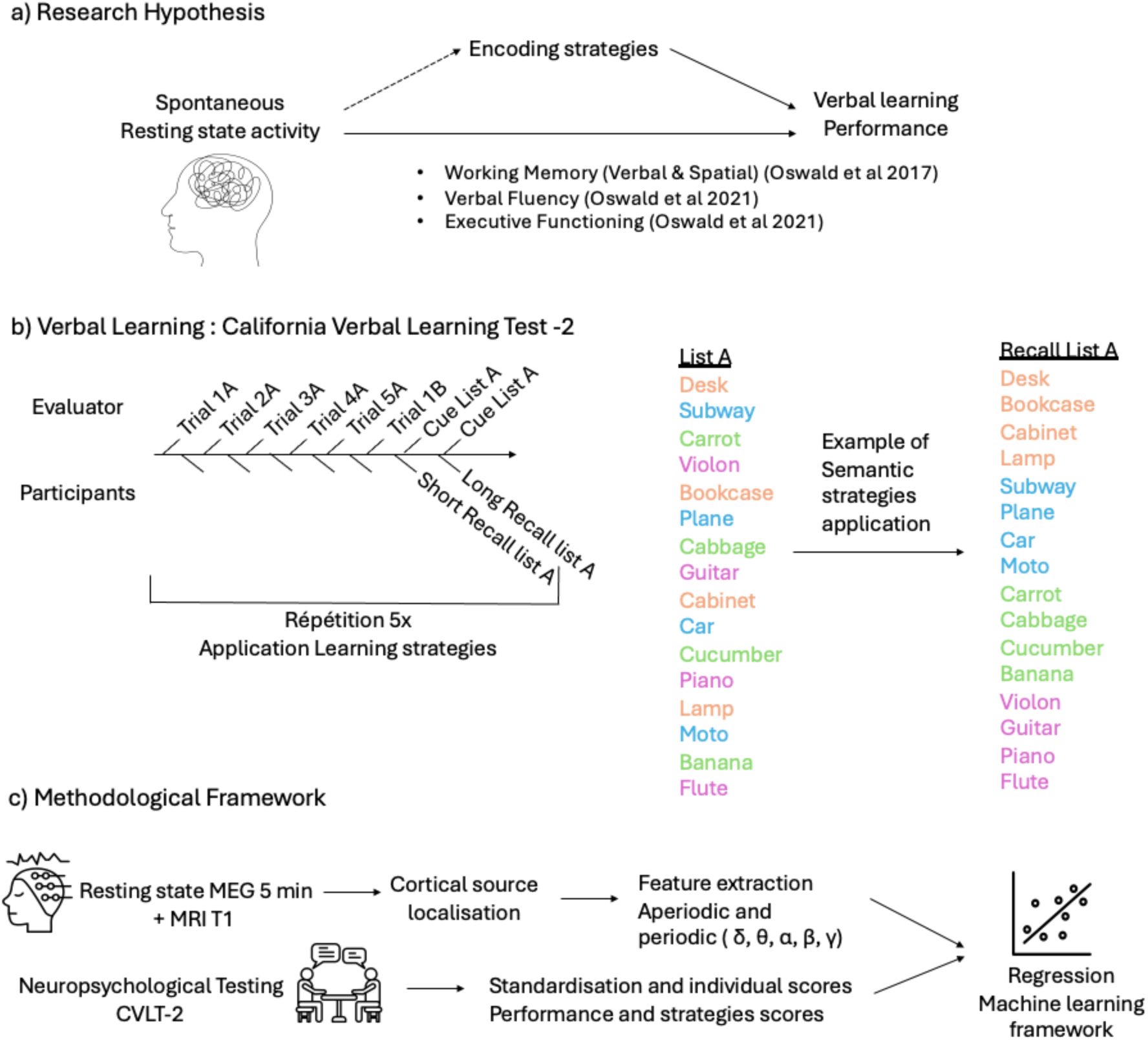
Global research framework, neuropsychological material, and methodological approach. **a)** Previous studies have investigated how spontaneous cognitive performance—such as working memory, verbal fluency, and executive functioning—is related to brain activity. Here, we test the hypothesis that the encoding strategies used during verbal learning tasks can be predicted by spontaneous neural activity. **b)** Example and explanatory view of the neuropsychological material: the California Verbal Learning Test–Second Edition (CVLT-II). **c)** Methodological framework: resting-state MEG data were collected prior to cognitive testing. Participants then completed a standardized neuropsychological assessment. Finally, a machine learning regression approach was applied across all subjects to predict individual encoding strategies from spontaneous brain activity.

The evaluation quantified the total number of correctly recalled words and the specific encoding strategies used by each participant. The CVLT-2 analysis included assessments of bidirectional (forward and backward) clustering strategies (counting the number of words recalled in the original or reverse order, respectively), semantic clustering (number of semantic groupings made within the four categories), and subjective clustering (number of two-words grouping made by a participant from the second trial and repeated in the next trial).

#### Correlations CVLT-2

To assess the benefits of each encoding strategy related to the performance memory, we regress the scores from the semantic, serial, and subjective strategies with overall performance with Pearson correlation method.

### Neuroimaging

#### Magnetoencephalography MEG and Anatomical MRI Data Acquisition

All 40 subjects were comfortably seated with eyes open, fixating on back-illuminated screen located 75cm in front of them. Two 5-minute periods of resting-state were recorded at a sampling rate of 1200Hz, using a CTF-VSM whole head 275-sensor MEG system (MEG core facility, Psychology Department, University of Montreal, QC, Canada). Following standard procedures, third-order gradiometer noise reduction was computed based on twenty-nine reference channels. Bipolar EOG (Vertical EOG and Horizontal EOG) was recorded to monitor eye blinks and ocular movements. ECG was also recorded to monitor heartbeats. Three head coils fixed at the nasion and the bilateral preauricular points were used for head localization and were monitored at the beginning and the end of each session. Care was taken to ensure that head displacement across sessions remained below 5 mm. The neuropsychological assessments were done in the morning at the Ste-Justine Hospital (Montreal, QC, Canada). Later in the afternoon, the participants went to the MEG facility, located in the Psychology Department of the University of Montreal, for the MEG recordings (Figure 1).

Structural MRI images were obtained for each subject with a 3-T General Electric (GE) scanner (Saint-Justine Hospital, Montreal, QC, Canada). The individual surfaces were used to carry out the co-registration between the MEG fiducial markers (LAP, NAS, RAP) and the MRI structural image. The exact position of the head was refined based on head shape position files obtained using a 3D-localization Polhemus system.

#### Data Pre-processing

MEG data pre-processing was performed using Brainstorm ^27^ – an open-source software based on MATLAB (Mathworks Inc., Natick, MA). The data was first notch filtered at 60Hz, and then between 0.5 Hz and 120 Hz. Cardiac artefacts, eye blinks, and eye movements were corrected using the Signal-Space-Projection method (SSP). Fifty signal epochs, centered on each artefact, were selected, and a singular value decomposition was applied to each artefact using built-in functions. Eigenvectors explaining at least 10 % of the variance of the artefacts were discarded and the remaining eigenvectors were used to define the SSP. The SSP method relies on a signal space decomposition procedure, where the statistical characteristics of the measured signals are used to determine the two subspaces spanned by the MEG brain signals and the unwanted artifacts, respectively. Projecting the continuous MEG data onto the signal subspace effectively removes the components belonging to the artifact subspace.

#### MEG Sources Estimation

MEG source reconstruction was performed using a standard weighted minimum-norm approach. T1-weighted brain volumes were acquired in all participants and were used to generate a cortical surface model using the FreeSurfer software package. Forward modeling of the magnetic field was defined based on an overlapping-sphere method. The weighted minimum norm solution was computed using a loose dipolar orientation constraint (set at 0.5), a signal-to-noise ratio of 3, whitening PCA and a depth weighting of 0.5. The noise covariance matrix for each participant was estimated from a 2-min empty room recording performed earlier the same day (same acquisition parameters but with no subject in the shielded room). The source time series were initially reconstructed on a 15000-vertex individual brain tessellation, and then spatially interpolated to the MNI ICBM152 brain template and down-sampled to a 10000-vertices template, and finally down-sampled to brain atlas of 300 resting state areas ^28^.

#### Spectral power analysis and correction for aperiodic component

The power spectrum density (PSD) was calculated using a modified Welch periodogram technique with a 3-second Hamming window and 50% time-window overlap at each cortical source. To disentangle and parameterize the periodic and aperiodic features of the PSD spectrum, the Fitting Oscillations and One-Over-F (FOOOF/spectrogram) algorithm, developed by ^29^, was employed. The FOOOF model was computed between 1Hz-120hz, with maximum peak at 3, detecting peak type with gaussian approach, proximity threshold at 2, and knee option for aperiodic mode selection. The 1/f model was subtracted from the full spectrum, and the frequencies were then averaged into specific frequency bands: Delta (1-4 Hz), theta (4-8 Hz), alpha (8-13 Hz), beta (13-30 Hz), gamma1 (30-60 Hz), gamma2 (60-90 Hz), gamma3 (90-120 Hz). Thus, we extracted the seven pure oscillatory components and then corrected for 1/f effect on oscillatory component effect.

#### Multi-feature machine learning regression analysis

To explore the relationship between aperiodic and oscillatory features and their impact on behavioral measures, we developed a multi-feature regression model using the Linear Support Vector Regressor (SVR) with 5-fold cross-validation. To optimize the model and improve its interpretability, we introduced sequential feature selection within a nested cross-validation framework. The goal was to identify the best combination of features, which we achieved through forward sequential feature selection with k-fold cross-validation (k=5). During each iteration, an individual feature is added to the model, starting with the most significant, and we assessed the model’s performance using metrics such as mean square error (MSE), r2, and p-values by comparing predicted data to the actual data. The subset of features that resulted in a highest significant r2 value (p < 0.05) was selected as the best set of features. To interpret the results and visualize the impact of the selected features, we extracted the coefficients associated with these features (ROIs) and weighted them by the data covariance to derive an activation function and to benefit the interpretation of regression outputs (Haufe et al., 2014). This approach provided insights into the relative importance of each selected feature in influencing behavioral learning strategies and learning performance measures. Our comprehensive approach aimed to identify the most relevant brain features contributing to the construction of the optimal brain model that fits the behavioral data. Additionally, for a large-scale perspective encompassing the entire brain, we also reported significant model coefficients for the entire brain and then most significant ROI. We are committed to transparency and collaboration; hence we provide and share the complete code used to compute this machine learning pipeline, enabling others to review, replicate, or build upon our work (https://github.com/LIKACT/CVLT_MEG_ML).

#### Statistical Analysis and Mediation analysis

To investigate the hypothesis that neural correlates are linked with learning strategies and learning performance, we employed a mediation model incorporating brain oscillatory ROIs as mediators. This model examined both single and serial dual mediator scenarios. Our approach began with a basic linear regression analysis to explore the relationships between various behavioral measures (including semantic clustering, serial clustering, subjective clustering, and learning performance scores) and individual brain ROIs across different frequencies. We selected only those ROIs demonstrating significant associations (p<0.05) as potential mediators in our subsequent analysis. The mediation models were then constructed and analyzed using the Hayes PROCESS toolbox. Distinct models were created for subjective and semantic clustering strategies. We applied the bootstrap method (n=5000) for testing both single mediator models and dual serial mediator models, focusing on identifying significant indirect pathways, defined as those showing a non-null range between the lower and upper confidence limits.

## Results

### Behavioral performance

Scores from the CVLT-2 were within the standard range (table 1). On average, participants recalled 58 word-items (std= 7.6) when cumulating the words recalled from first trial to the fifth trial with the list A. Table 1 shows the mean performance across the five trials, and the mean clustering for every learning strategy, namely semantic, serial bidirectional and subjective. Our results showed that participants predominantly favored the semantic strategy (mean=1.95, std =2.62), followed by the subjective clustering strategy (mean=1.53, std =1.17) and then by the bidirectional serial strategy (mean = 0.90, std=1.78).

**Table 1:**
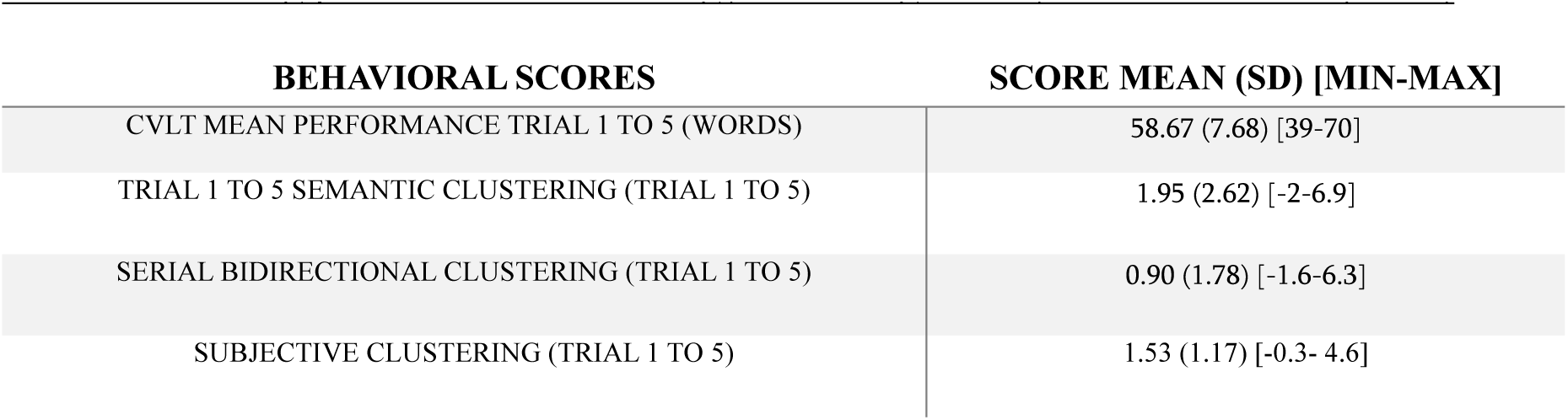
Learning performance and strategy clustering trial by trial in CVLT-2 (n=40)

Figure 2 shows linear regressions between mean raw performance across trials 1 to 5 and mean indices of each strategy used across the five trials. Result show that mean learning performance is strongly predicted by subjective clustering (r2= .38; p<.001), as well as by semantic clustering (r2= .21; p =.003), while bidirectional serial clustering did not significatively regress with mean performance (r2=0; p =.91). The confusion matrix (figure 2d) shows the strong anticorrelation between semantic clustering and serial clustering (r=-0.8; p>0.001), while correlation between semantic clustering and subjective clustering were found significant (r=0.32; p=0.04). No significant relationship between serial clustering and subjective clustering was found (r=0.21; p=0.18).

**Figure 2:**
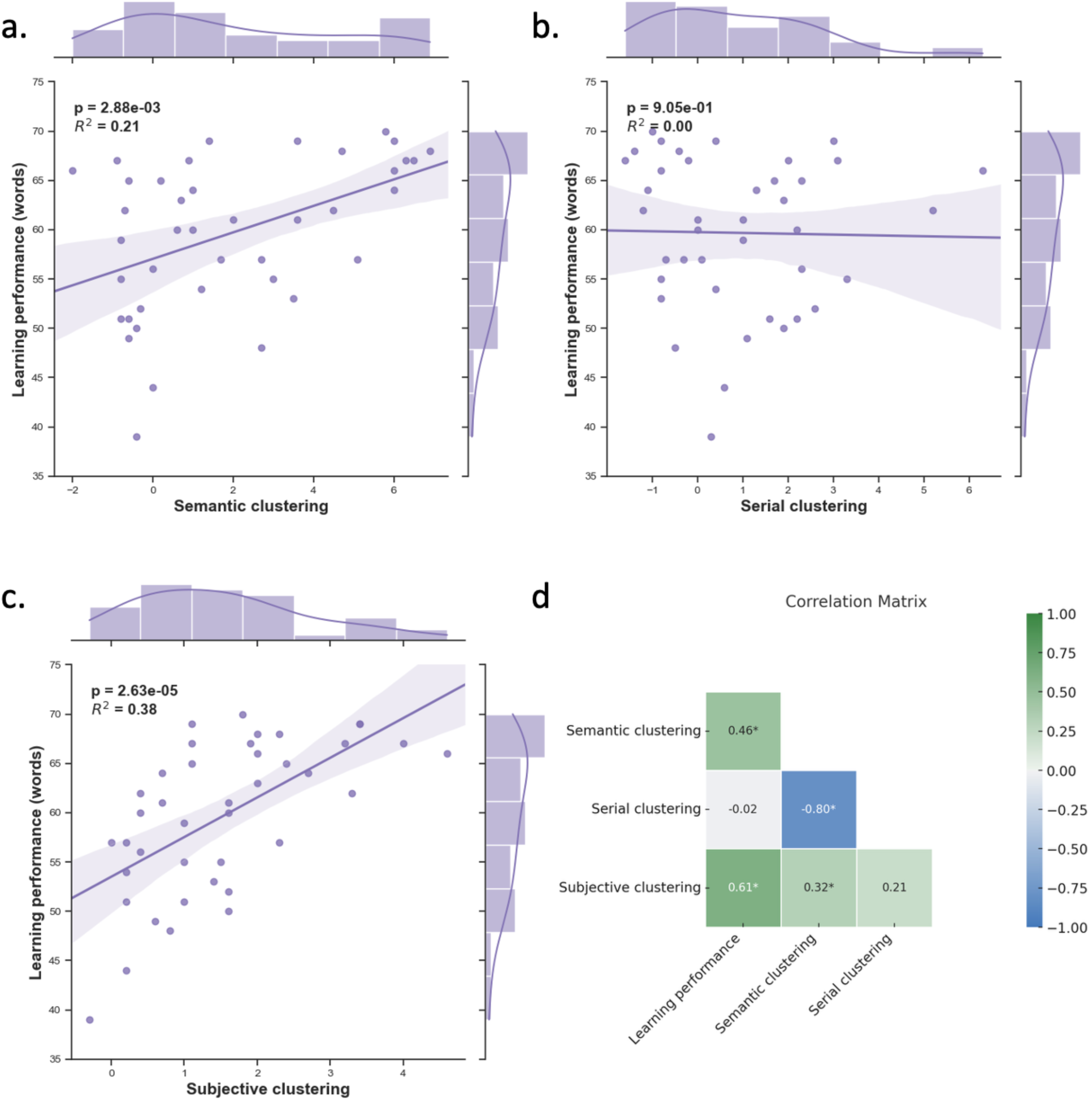
Behavioral relationship between verbal learning performance and encoding strategies, a) Linear regression between mean performance (words) and semantic clustering; b) Linear regression between mean performance (words) and serial clustering; c) Linear regression between mean performance (words) and subjective clustering; d) matrix confusion correlation between verbal learning performance and all encoding strategies.

### Brain Behavior for verbal learning performance

#### Verbal learning performance

The results of our machine learning model across the whole brain demonstrate that resting-state MEG theta power could predict the performance of the number of words learned during the CVLT-2 test. The model for the entire brain showed an explained variance of r2 = 0.16 (p = 0.01) (Figure 3), while the best result obtained with feature selection (ROI) used 93 out of 300 ROIs, with an r2=0.2 (p = 0.003) (Figure S1). The distribution of weights across the brain demonstrated a positive association between theta oscillations and the sensorimotor and parietal regions of the left hemisphere, as well as bilaterally in the premotor regions. Conversely, bilateral visual and limbic regions were strongly negatively associated with verbal learning performance to theta oscillations.

**Figure 3:**
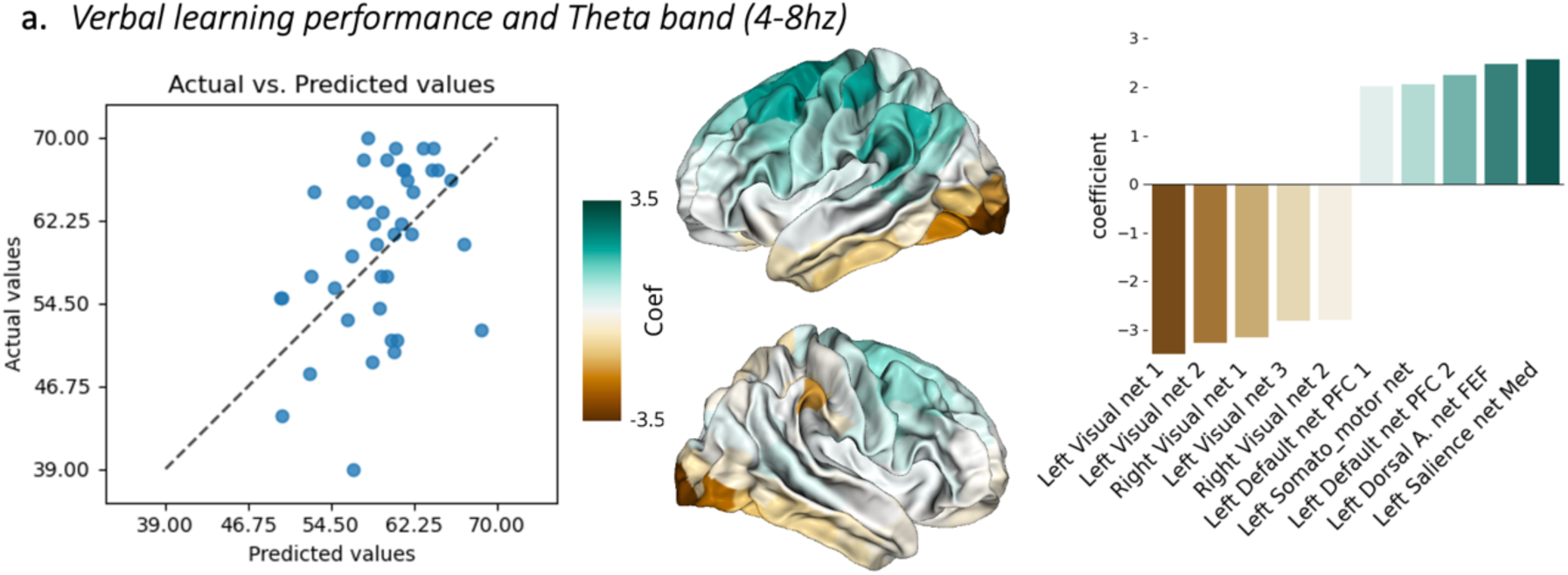
Brain-behavior machine learning regression (linear SVR) between multi-feature (brain regions) resting state oscillatory components and the **correct responses scores.** This includes a brain map projection of the regression coefficients associated with data covariance (left), and the average coefficient across the resting state network atlas (right side)

### Brain Behavior for encoding strategies

#### Subjective clustering

The subjective encoding strategy, which involves recalling two different elements of a word list in the same or inverse order they were recalled on the immediate preceding trial, showed a significant regression model with r2 = 0.09 (p = 0.04) between resting theta power and the average score of employing this encoding strategy during the learning of the word list (Figure 4). The distribution of weights across the brain revealed a positive association between bilateral premotor, sensorimotor and parietal regions, with the largest values in the left hemisphere. Conversely, bilateral orbito-prefrontal and inferior and medial temporal regions (limbic network) were strongly negatively associated with subjective clustering.

**Figure 4:**
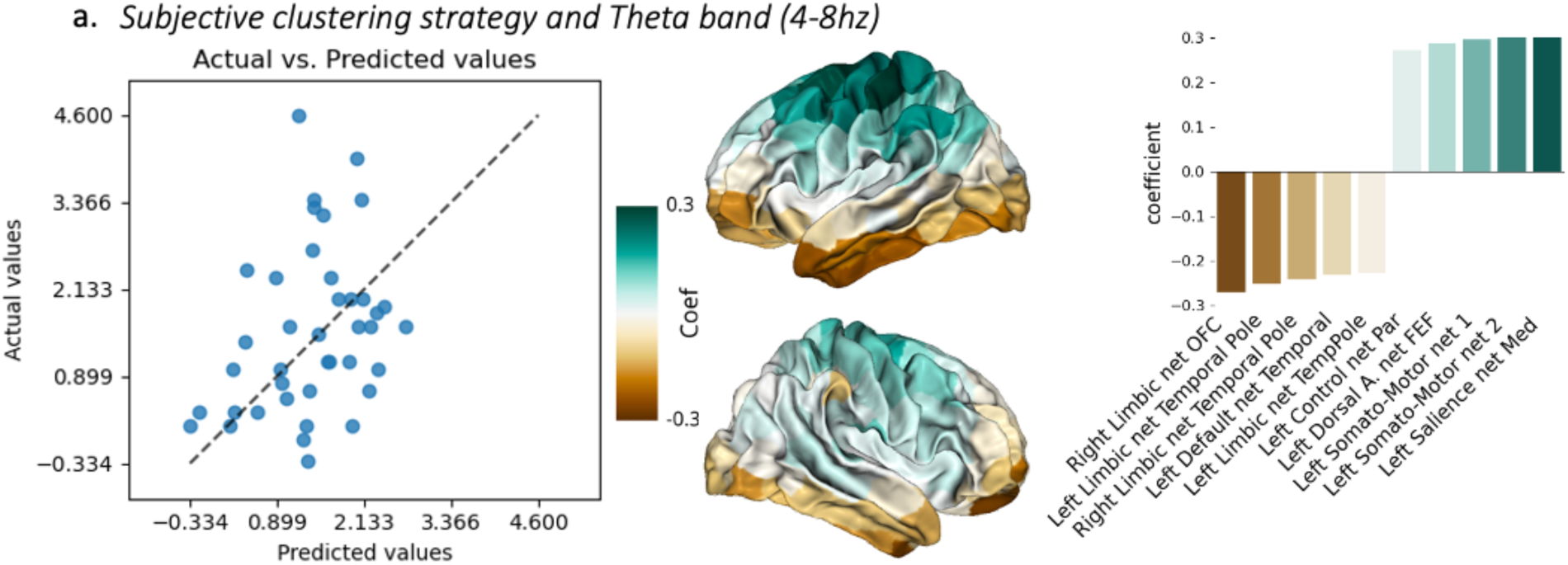
Brain-behavior machine learning regression (linear SVR) between multi-feature (brain regions) resting state oscillatory components and the **subjective clustering learning strategy**. This includes a brain map projection of the regression coefficients associated with data covariance (left), and the average coefficient across the resting state network atlas (right side)

#### Semantic clustering

The encoding strategy scores using semantic clustering, which involves grouping different elements of a word list into semantic categories to facilitate memorization, demonstrated a significant association with resting alpha power (r2 = 0.13; p = 0.01), beta power (r2 = 0.13; p = 0.01), and gamma (60-90 Hz) power (r2 = 0.09; p = 0.05) (Figure 5). The best feature-selected model in the same frequency bands, respectively, were alpha with 93 out of 300 ROIs (r2 = 0.15; p = 0.01); 58 out of 300 ROIs for the beta band (r2 = 0.22; p = 0.001); and 215 out of 300 ROIs for gamma (60-90 Hz) (r2 = 0.13; p = 0.02) (Figure S1). The distribution of weights across the brain for the alpha and beta bands demonstrated a strong left lateralization with a positive association between resting power and the use of the semantic strategy in the sensorimotor and parietal regions, including the superior part of the left temporal lobe (somato-motor and control network). Conversely, a negative association was found with the right visual regions as well as the lower temporal regions (limbic and visual network). For the gamma band, the same spatial distribution was found, but the direction of the association between oscillations and behavior is reversed.

**Figure 5:**
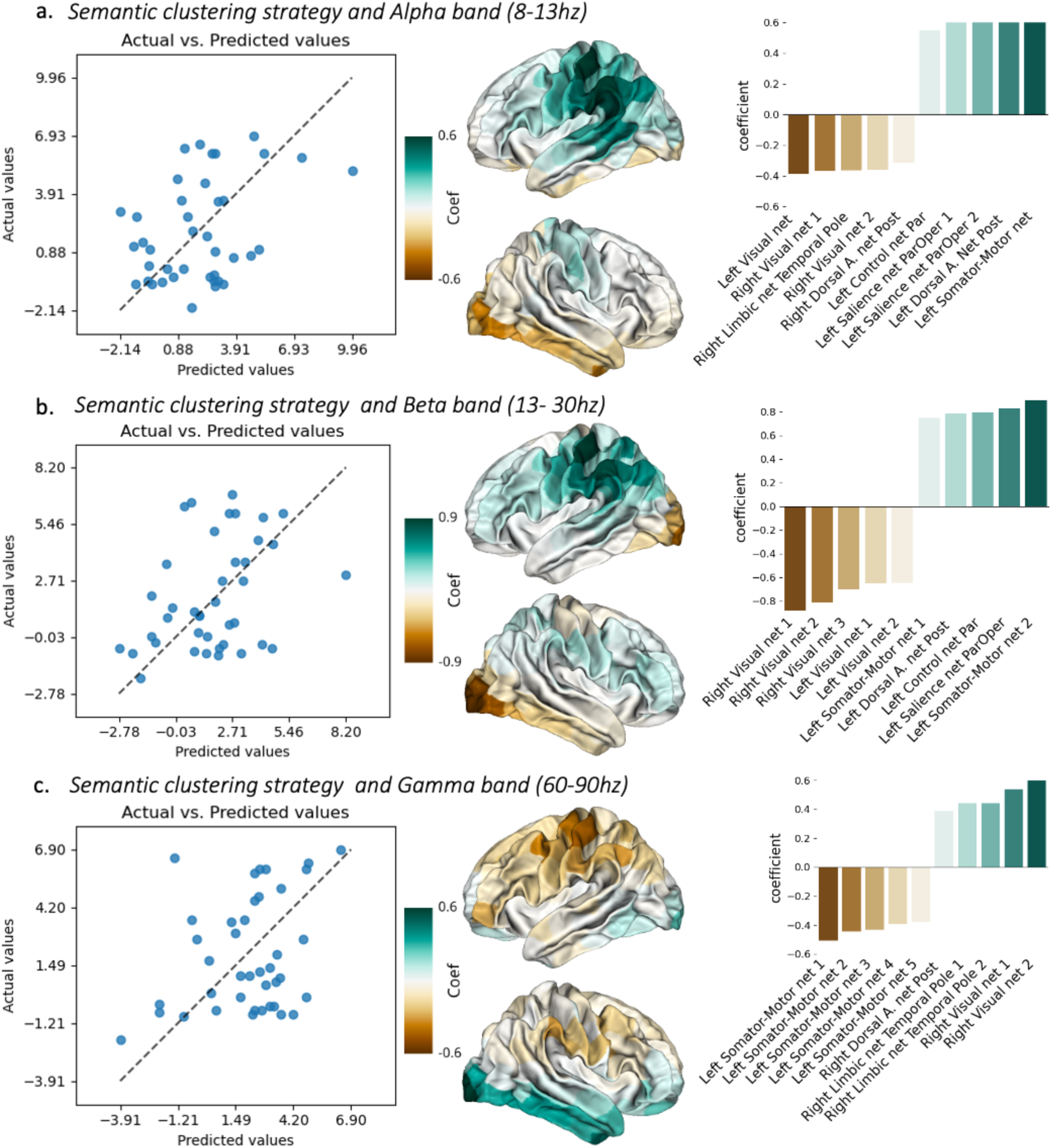
Brain-behavior machine learning regression (linear SVR) between multi-feature (brain regions) resting state oscillatory components and the **semantic clustering learning strategy**. This includes a brain map projection of the regression coefficients associated with data covariance (left), and the average coefficient across the resting state network atlas (right side)

#### Serial bilateral clustering

The encoding strategy scores using serial bidirectional clustering, which involves grouping items in the same or inverse order as they were presented during the word list, demonstrated a significant association with resting beta power (r2 = 0.21; p = 0.002) and gamma (60-90 Hz) power (r2 = 0.12; p = 0.02) (Figure 6). The best feature-selected model in the same frequency bands, respectively, used 213 out of 300 ROIs for the beta band (r2 = 0.24; p = 0.001); and 234 out of 300 ROIs for gamma (60-90 Hz) (r2 = 0.15; p = 0.01) (Figure S1). The distribution of weights across the brain for the beta band revealed a left lateralization with a negative association between resting power and the use of the serial strategy in sensorimotor and parietal regions, including the temporo-occipital regions (control and salience network). Conversely, a positive association was observed with the right dorsal sensorimotor cortex as well as the lower right temporal regions (limbic and somato-motor network). For the gamma band, bilateral implications of dorsal regions were found to positively correlate between the gamma band and serial clustering (somato-motor and salience network). The strongest effect was found in the left hemisphere compared to the right hemisphere. Conversely, a strong negative association was detected with the lower part of the right temporal lobe as well as the right prefrontal cortex (limbic and visual network).

**Figure 6:**
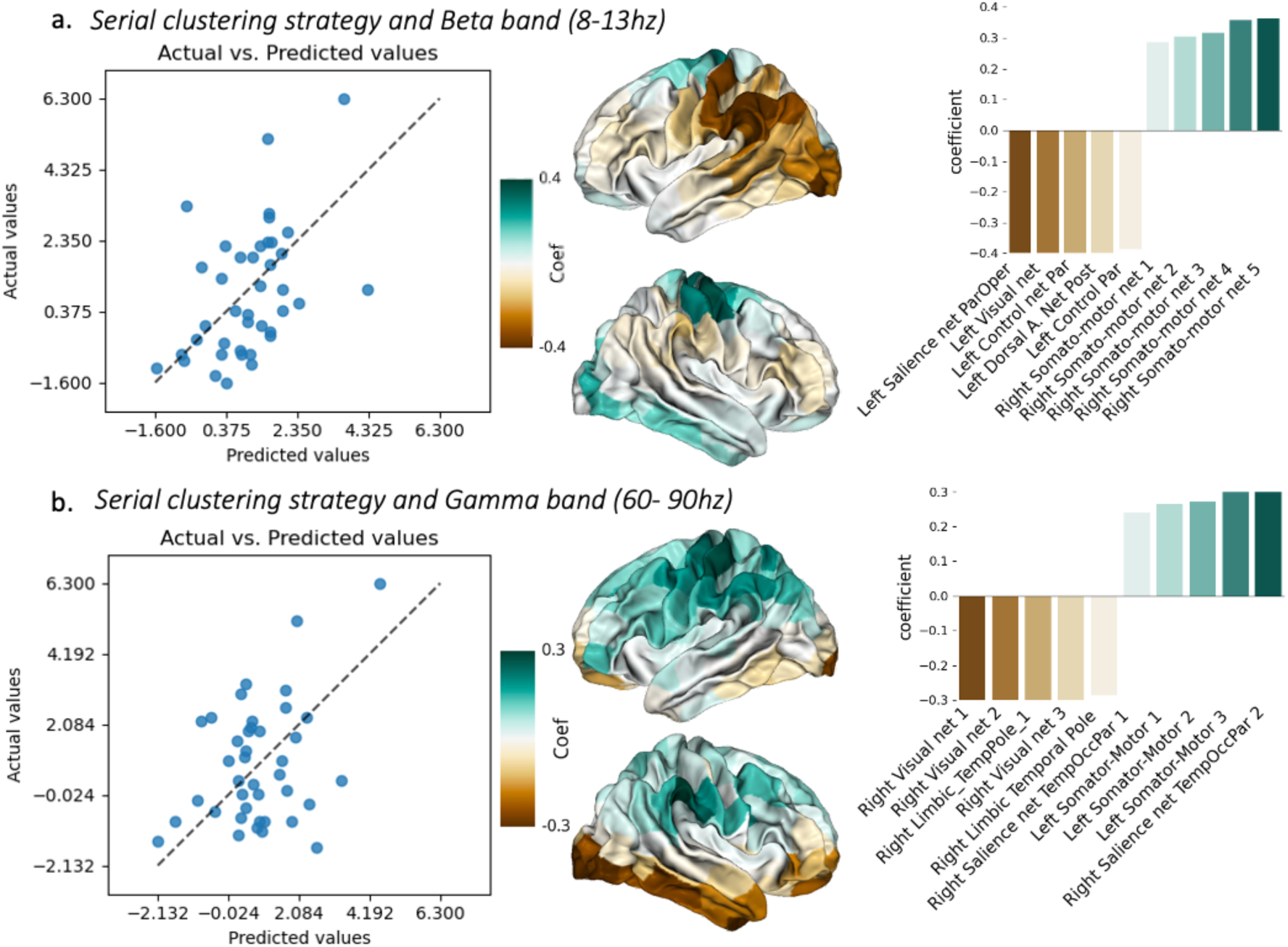
Brain-behavior machine learning regression (linear SVR) between multi-feature (brain regions) resting state oscillatory components and the **serial clustering learning strategy**. This includes a brain map projection of the regression coefficients associated with data covariance (left), and the average coefficient across the resting state network atlas (right side)

### Opposing Spatio-Frequency Patterns Associated with Serial and Semantic Strategies

To highlight the main results emerging from the different frequency-spatial-strategy patterns, we summarized the information in Figure 7. This figure aims to illustrate the different oppositions based on the weight of the coefficients derived from the multi-feature models. For example, beta oscillations are strongly and positively predictive of semantic strategy use in certain brain regions (e.g., control and somato-motor networks), meaning that higher resting-state beta activity predicts greater semantic clustering. Conversely, the same regions show a negative association between beta power and serial strategy use (i.e., higher beta at rest predicts reduced serial clustering). These opposing relationships are primarily observed in the control (fronto-parietal), somato-motor, visual, and limbic networks. Moreover, similar oppositions are seen in the associations between beta and gamma-band activity (60–90 Hz) with semantic and serial strategies. In summary, at rest, a dual opposition — both oscillatory and spatial — characterizes the alternate recruitment of semantic and serial memory strategies.

**Figure 7:**
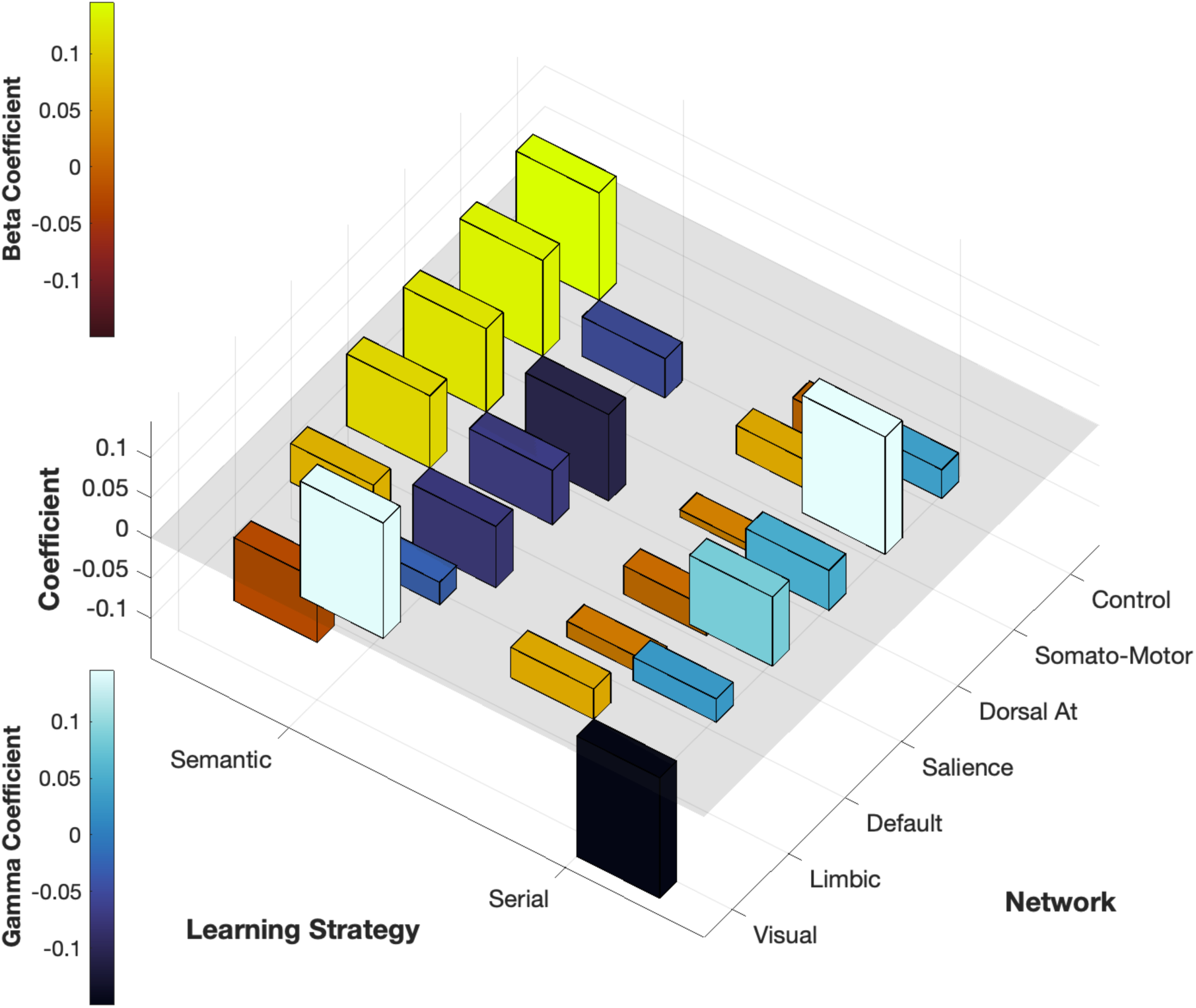
Summary of Brain–Behavior Associations for Semantic and Serial Strategies Across Beta and Gamma Bands (60–90 Hz). Spatial maps display the direction and strength of associations between resting-state oscillatory activity and strategies across brain regions. Light colors indicate positive associations (e.g., higher beta predicting greater semantic clustering), while dark colors indicate negative associations (e.g., higher beta predicting reduced serial clustering). Opposing patterns are observed across control, somato-motor, visual, and limbic networks, highlighting distinct spatial and frequency-specific recruitment for semantic vs. serial strategies.

## Discussion

In this study, we investigated the relationship between spontaneous neural oscillations and verbal learning performance, along with the associated encoding strategies, by combining resting-state MEG with a standardized neuropsychological memory assessment (CVLT-2). Overall, group performance was within the expected normative range for this test ^23^. Both semantic and subjective clustering strategies were significantly correlated with overall learning performance (Figure 2), whereas the serial strategy showed no such association. Moreover, we observed shared variance between semantic and subjective clustering strategies, with subjective clustering emerging as the strongest predictor of performance. The CVLT-2 design inherently permits the concurrent use of both strategies. Although the CVLT-2 cannot fully disentangle clustering strategies, it benefits from not enforcing any particular one. Beyond its verbal nature and the potential for semantic or serial organization, the lack of explicit strategic instruction means that strategy use remains largely spontaneous, reflecting individual cognitive tendencies rather than guided behavior ^24^.

### Resting-state patterns support encoding strategies

Our neuroimaging results revealed distinct spatio-frequency patterns that differentiate not only overall learning performance but also the spontaneous use of subjective, semantic, and serial clustering strategies. After removing the 1/f aperiodic component from the power spectral density, we found that learning performance and subjective clustering were primarily associated with resting-state theta oscillations (figure 3 & 4). In contrast, semantic and serial clustering strategies were associated with alpha, beta, and gamma oscillatory activity (figure 5 & 6). At the spatial level, learning performance was negatively associated with bilateral visual regions and positively associated with left parietal, sensorimotor, and bilateral premotor cortices. In turn, the subjective clustering strategy was negatively associated with bilateral occipital, inferior, and medial temporal regions, and positively associated with left-lateralized premotor, sensorimotor, and parietal regions. Across both the alpha and beta bands, both semantic and serial clustering strategies were positively associated with posterior sensorimotor regions and negatively associated with posterior temporal, visual, and inferior temporal areas. These associations reversed in the gamma band, suggesting frequency-specific tuning for these strategies.

Together, these findings highlight that resting-state neural activity is not only related to individual differences in learning performance but also reflects the spontaneous cognitive strategies adopted during memory encoding. This aligns with prior studies showing that resting-state oscillatory activity can predict individual cognitive styles, such as a preference for analytical versus insight-based problem-solving ^31^, as well as working memory efficiency and attention control ^32^. Moreover, recent evidence suggests that the dynamic variability and spectral characteristics of resting-state networks are robust predictors of interindividual differences in higher-order cognition, including encoding-related processes^33^. To our knowledge, this is the first study to demonstrate that spontaneous spatio-frequency resting-state patterns can distinguish between distinct verbal encoding strategies, suggesting that intrinsic neural architecture supports not only memory capacity but also strategic processing styles.

### Learning and Subjective encoding strategies

We observed similarities in both spatial and frequency patterns between the subjective clustering strategy and learning performance, suggesting a shared underlying mechanism centered on theta oscillations. These similarities indicate that an individual’s memory performance is closely tied to their preferred recall strategy. Given their shared variance (figure 2), a substantial proportion of recall performance may be explained by idiosyncratic encoding strategies that are neurally instantiated within the brain’s resting-state functional organization.

To our knowledge, this is among the first demonstrations that spontaneous neuronal organization at rest encodes interindividual variability in encoding strategies. While previous research has investigated this issue primarily through task-based neuroimaging paradigms, numerous studies have shown that the use of self-initiated encoding strategies influences memory performance and is associated with specific brain regions. For example, functional imaging studies have linked strategic encoding to activity in the prefrontal cortex ^34^, the left medial temporal lobe ^35^ and the left lingual gyrus ^36^, as well as to networks encompassing the left fusiform gyrus, right cingulate cortex, right posterior hippocampus, bilateral retrosplenial cortex, and left medial superior parietal regions, depending on the type of strategy employed ^37–39^; see review - (Kirchhoff, 2009). Our resting-state findings are consistent with this literature, showing the involvement of medial and inferior temporal regions (including parahippocampal, lingual, and fusiform gyri) and prefrontal regions in the subjective clustering strategy. Importantly, theta oscillations—well known to support memory encoding during active tasks ^41,42^ and to scale with memory load (e.g., Raghavachari et al., 2001) were here predictive of subsequent learning performance at rest, via the recruitment of subjective clustering. Given that subjective strategies rely on repeating the same recall pattern across trials, we propose that resting-state theta activity reflects a stable, individual-specific mode of associative learning. This interpretation aligns with converging evidence from human studies showing that theta oscillations are central to both encoding and retrieval processes ^44^. In extending prior task-based findings, our results show that resting-state neural patterns not only capture these effects at the oscillatory level but also reveal their corresponding spatial substrates in the brain.

### Semantic and Serial encoding strategies

A second major focus of our study concerns the semantic and serial encoding strategies. These strategies are discussed together because they are behaviorally related— often showing anticorrelation at the individual level (Figure 2)—and because they are supported by opposite, yet clearly defined spatio-frequency patterns (figure 7). Both are primarily sustained by oscillatory activity in the alpha, beta, and gamma bands, although their spatial distributions and functional interpretations diverge.

The semantic clustering strategy was associated with a left-lateralized spatial pattern encompassing the premotor cortex, postcentral and precentral gyri, superior and inferior parietal lobules, superior temporal gyrus, and posterior middle and inferior temporal gyri. This extensive network is consistent with the well-established literature on semantic memory, which highlights a distributed neural architecture supporting multiple cognitive components: Language processing — the left superior temporal sulcus and left inferior lateral frontal gyrus are critical for linguistic analysis ^45–47^. Conceptual and knowledge representation — the middle, anterior, and posterior temporal gyri serve as core hubs for conceptual knowledge and semantic integration ^48–50^. Motor planning and action representation — the premotor, motor, and somatosensory cortices contribute to the embodied representation of semantic concepts ^51,52^. From an embodied cognition perspective, semantic representations are partly grounded in sensory–motor cortices, with conceptual knowledge encoded in regions linked to action-related motor representations ^53–55^. In the CVLT-2 paradigm used here, the 16 words are organized into four semantic categories (vegetables, furniture, animals, transportation). Although words from the same category are never presented consecutively, participants can rapidly detect the categorical structure and apply a category-based abstract representation for learning. These categories align with the types of information classically encoded by semantic memory ^53–55^.

Oscillatory analysis revealed that semantic clustering was supported by alpha, beta, and gamma activity. Alpha and beta oscillations are extensively studied in the context of short-term and working memory, particularly within fronto-parietal networks ^7,56–59^. These rhythms are thought to implement functional inhibition, allowing for more efficient manipulation and organization of information ^60–63^. In a verbal learning context, this inhibitory gating may facilitate the mental grouping of words into semantic categories (as shown in Figure 1). Previous task-based studies have shown clear relationships between semantic processing and alpha ^47,64–66^, beta ^10,67,68^, and gamma oscillations ^69^. Our findings extend this literature by showing that spontaneous resting-state activity in these frequency bands predicts the subsequent use of semantic strategies, suggesting that the neural basis for semantic organization is partly intrinsic and observable even in the absence of an explicit task.

In contrast to semantic clustering—which engaged a left-lateralized temporal– parietal–frontal network—the serial clustering strategy involved a more fronto-parietal and motor-planning profile, including the dorsolateral prefrontal cortex, supplementary motor area, precuneus, and posterior parietal regions. These areas are consistently linked to maintaining and manipulating item order in working memory ^70,71^ and to temporal structuring of recall ^72^. Like semantic clustering, serial clustering was supported by beta and gamma oscillations, but with opposite associations in key overlapping regions (Figure 7). Such patterns align with evidence that beta suppression and gamma enhancement in fronto-parietal circuits could facilitate sequential order processing ^73^ and support the view that these strategies compete for shared neural resources.

One of the major clarifications provided by our results is the frequency-specific distinction between encoding strategies. Two primary effects emerged: Theta vs. Beta/Gamma separation — subjective clustering (discussed earlier) was predominantly supported by theta oscillations, whereas semantic and serial clustering strategies were supported by beta and gamma oscillations. Opposite beta/gamma patterns in the same regions — regions that showed a positive association with beta and gamma power for semantic clustering exhibited negative associations with these same bands associate with serial clustering. This reversal suggests that resting-state spectral power in specific cortical regions predisposes individuals to favor either a semantic or a serial clustering approach when encoding new information. The presence of opposing frequency signatures within the same anatomical regions supports the idea of functional competition: adopting one strategy may suppress the oscillatory dynamics necessary for the other, explaining their behavioral anticorrelation^74^. To our knowledge, this is the first study to demonstrate that spontaneous oscillatory patterns at rest can predict the preferential use of semantic versus serial encoding strategies, underscoring the capacity of intrinsic brain dynamics to shape individual cognitive styles

### Limits and Perspectives

Replication of these findings in larger datasets, whether using electrophysiology or large-scale fMRI, would provide stronger validation and generalizability. Confirming these results could lead to two major applications. First, it would open the possibility of targeted strategy training—for example, explicitly teaching participants to use semantic clustering and assessing whether this training modulates oscillatory activity (e.g., alpha or beta) both during tasks and at rest. Second, given the CVLT-2’s recruitment of diverse cognitive processes, and its established role as a diagnostic tool in clinical neuropsychology for various psychopathologies, understanding its neural underpinnings could clarify the neurophysiological systems disrupted in these disorders. If strategy training is shown to induce measurable neural changes, this approach could inform therapeutic interventions or neuromodulation protocols aimed at enhancing cognitive function via the modulation of underlying oscillatory dynamics.

### Conclusion

This study provides the first direct evidence linking resting-state cortical spectral power, as measured with MEG, to the spontaneous adoption of verbal learning strategies assessed with a standardized neuropsychological tool (CVLT-2) outside the scanner. These findings advance our understanding of the neuronal mechanisms supporting encoding strategies and highlight the role of intrinsic brain dynamics in shaping cognitive styles. By bridging neurophysiological signatures at rest with behavioral strategy use, this work opens new avenues for both basic research and potential clinical applications.

## References

1. Florin, E. & Baillet, S. The brain’s resting-state activity is shaped by synchronized cross-frequency coupling of neural oscillations. Neuroimage 111, 26–35 (2015).

2. Brookes, M. J. et al. Investigating the electrophysiological basis of resting state networks using magnetoencephalography. Proc Natl Acad Sci U S A 108, 16783– 16788 (2011).

3. Buzsaki, G. & Draguhn, A. Neuronal Oscillations in Cortical Networks. Science (1979) 304, 1926–1929 (2004).

4. Ossandón, T. et al. Transient suppression of broadband gamma power in the default-mode network is correlated with task complexity and subject performance. Journal of Neuroscience 31, 14521–14530 (2011).

5. Palva, S. & Palva, J. M. New vistas for α-frequency band oscillations. Trends Neurosci 30, 150–158 (2007).

6. Jerbi, K. et al. Exploring the electrophysiological correlates of the default-mode network with intracerebral EEG. Front Syst Neurosci 4, 1–9 (2010).

7. Oswald, V. et al. Spontaneous brain oscillations as neural fingerprints of working memory capacities: A resting-state MEG study. Cortex 97, 109–124 (2017).

8. Clark, C. R. et al. Spontaneous alpha peak frequency predicts working memory performance across the age span. International Journal of Psychophysiology 53, 1–9 (2004).

9. Riddle, J., Scimeca, J. M., Cellier, D., Dhanani, S. & D’Esposito, M. Causal Evidence for a Role of Theta and Alpha Oscillations in the Control of Working Memory. Current Biology 30, 1748–1754.e4 (2020).

10. Oswald, V. et al. Magnetoencephalography resting-state correlates of executive and language components of verbal fluency. Sci Rep 12, 1–12 (2022).

11. Baddeley, A. The episodic buffer: a new component of working memory? Trends Cogn Sci 4, 6267–6279 (2000).

12. Baddeley, A. D. & Hitch, G. Working Memory. in (ed. Bower, G. H. B. T.-P. of L. and M.) vol. 8 47–89 (Academic Press, 1974).

13. Tulving, E. Episodic and semantic memory. in Organization of memory. xiii, 423– xiii, 423 (Academic Press, Oxford, England, 1972).

14. Hertzog, C., Price, J. & Dunlosky, J. How is knowledge generated about memory encoding strategy effectiveness? Learn Individ Differ 18, 430–445 (2008).

15. Miotto, E. C. et al. Bilateral activation of the prefrontal cortex after strategic semantic cognitive training. Hum Brain Mapp 27, 288–295 (2006).

16. Miotto, E. C. et al. Effects of semantic categorization strategy training on episodic memory in children and adolescents. PLoS One 15, 1–17 (2020).

17. Miotto, E. C. et al. Brain regions supporting verbal memory improvement in healthy older subjects. Arq Neuropsiquiatr 72, 663–670 (2014).

18. Kirchhoff, B., Anderson, B., Barch, D. & Jacoby, L. Cognitive and neural effects of semantic encoding strategy training in older adults. Cerebral Cortex 22, 788– 799 (2012).

19. Turner, A. D., Furey, M. L., Drevets, W. C., Zarate, C. & Nugent, A. C. Association between subcortical volumes and verbal memory in unmedicated depressed patients and healthy controls. Neuropsychologia 50, 2348–2355 (2012).

20. Baldo, J. V, Delis, D., Kramer, J. & Shimamura, A. P. Memory performance on the California Verbal Learning Test – II : Findings from patients with focal frontal lesions. 539–546 (2020).

21. Baldo, J. V., SChwartz, S., Wilkins, D. & Dronkers, N. F. Role of frontal versus temporal cortex in verbal fluency as revealed by voxel-based lesion symptom mapping. Journal of the International Neuropsychological Society 12, 896–900 (2006).

22. Gongvatana, A., Woods, S. P., Taylor, M. J., Vigil, O. & Grant, I. Semantic clustering inefficiency in HIV-associated dementia. Journal of Neuropsychiatry and Clinical Neurosciences 19, 36–42 (2007).

23. Elwood, R. W. The California Verbal Learning Test: Psychometric characteristics and clinical application. Neuropsychol Rev 5, 173–201 (1995).

24. Thiruselvam, I. & Hoelzle, J. B. Refined Measurement of Verbal Learning and Memory: Application of Item Response Theory to California Verbal Learning Test - Second Edition (CVLT-II) Learning Trials. Arch Clin Neuropsychol 35, 90–104 (2020).

25. Bradley, V. & Kapur, N. Neuropsychological assessment of memory disorders. in University Press Scholarship Online Oxford Scholarship Online (2010). doi:10.1093/acprof.

26. D.C. Delis, J.H. Kramer, E. Kaplan, B. A. O. California Verbal Learning Test – second edition. Adult version. Manual. *Psychological Corporation,San Antonio*, TX (2000).

27. Tadel, F., Baillet, S., Mosher, J. C., Pantazis, D. & Leahy, R. M. Brainstorm: A user-friendly application for MEG/EEG analysis. Comput Intell Neurosci 2011, (2011).

28. Schaefer, A. et al. Local-Global Parcellation of the Human Cerebral Cortex from Intrinsic Functional Connectivity MRI. Cerebral Cortex 28, 3095–3114 (2018).

29. Donoghue, T. et al. Parameterizing neural power spectra into periodic and aperiodic components. Nat Neurosci 23, 1655–1665 (2020).

30. Haufe, S. et al. On the interpretation of weight vectors of linear models in multivariate neuroimaging. Neuroimage 87, 96–110 (2014).

31. Erickson, B. et al. Resting-state brain oscillations predict trait-like cognitive styles. Neuropsychologia 120, 1–8 (2018).

32. Sargent, K. et al. Resting-state brain oscillations predict cognitive function in psychiatric disorders: A transdiagnostic machine learning approach. Neuroimage Clin 30, (2021).

33. Chen, C. et al. Resting-state EEG network variability predicts individual working memory behavior. Neuroimage 310, (2025).

34. Savage, C. R. et al. Prefrontal regions supporting spontaneous and directed application of verbal learning strategies. Brain 124, 219–231 (2001).

35. Frings, L. et al. Gender-related differences in lateralization of hippocampal activation and cognitive strategy. Neuroreport 17, 417–421 (2006).

36. Heinze, S. et al. Neural encoding correlates of high and low verbal memory performance. J Psychophysiol 20, 68–78 (2006).

37. Nohara, S. et al. Neural correlates of memory organization deficits in schizophrenia: A single photon emission computed tomography study with 99mTc-ethyl-cysteinate dimer during a verbal learning task. Schizophr Res 42, 209–222 (2000).

38. Hazlett, E. A. et al. Hypofrontality in unmedicated schizophrenia patients studied with PET during performance of a serial verbal learning task. Schizophr Res 43, 33–46 (2000).

39. Maguire, E. A., Valentine, E. R., Wilding, J. M. & Kapur, N. Routes to remembering: The brains behind superior memory. Nat Neurosci 6, 90–95 (2003).

40. Kirchhoff, B. A. Individual differences in episodic memory: The role of self-initiated encoding strategies. Neuroscientist 15, 166–179 (2009).

41. ter Wal, M., et al. Theta rhythmicity governs human behavior and hippocampal signals during memory-dependent tasks. Nat Commun 12, 1–15 (2021).

42. Bastiaansen, M. C. M., Van Der Linden, M., Ter Keurs, M., Dijkstra, T. & Hagoort, P. Theta responses are involved in lexical-semantic retrieval during language processing. J Cogn Neurosci 17, 530–541 (2005).

43. Raghavachari, S., et al. Gating of Human Theta Oscillations by a Working Memory Task. (2001).

44. Herweg, N. A., Solomon, E. A. & Kahana, M. J. Theta Oscillations in Human Memory. Trends Cogn Sci 24, 208–227 (2020).

45. Halgren, E. et al. N400-like magnetoencephalography responses modulated by semantic context, word frequency, and lexical class in sentences. Neuroimage 17, 1101–1116 (2002).

46. Lopes, T. M. et al. Effects of task complexity on activation of language areas in a semantic decision fMRI protocol. Neuropsychologia 81, 140–148 (2016).

47. Mousavi, N., Nazari, M. A., Babapour, J. & Jahan, A. Electroencephalographic characteristics of word finding during phonological and semantic verbal fluency tasks. Neuropsychopharmacol Rep 40, 254–261 (2020).

48. Elger, C. E. et al. Human temporal lobe potentials in verbal learning and memory processes. Neuropsychologia 35, 657–667 (1997).

49. Kapur, S. et al. The neural correlates of intentional learning of verbal materials: A PET study in humans. Cognitive Brain Research 4, 243–249 (1996).

50. Saykin, A. J. et al. Functional differentiation of medial temporal and frontal regions involved in processing novel and familiar words: An fMRI study. Brain 122, 1963–1971 (1999).

51. Binder, J. R. & Desai, R. H. The neurobiology of semantic memory. Trends Cogn Sci 15, 527–536 (2011).

52. Binder, J. R., Desai, R. H., Graves, W. W. & Conant, L. L. Where is the semantic system? A critical review and meta-analysis of 120 functional neuroimaging studies. Cerebral Cortex 19, 2767–2796 (2009).

53. Gallese, V. & Lakoff, G. The brain’s concepts: The role of the sensory-motor system in conceptual knowledge. Cogn Neuropsychol 22, 455–479 (2005).

54. Gallese, V. & Cuccio, V. The neural exploitation hypothesis and its implications for an embodied approach to language and cognition: Insights from the study of action verbs processing and motor disorders in Parkinson’s disease. Cortex 100, 215–225 (2018).

55. Binder, J. R. et al. Toward a brain-based componential semantic representation. Cogn Neuropsychol 33, 130–174 (2016).

56. Mamashli, F., Khan, S., Obleser, J., Friederici, A. D. & Maess, B. Oscillatory dynamics of cortical functional connections in semantic prediction. Hum Brain Mapp 40, 1856–1866 (2019).

57. Klimesch, W. Alpha-band oscillations, attention, and controlled access to stored information. Trends Cogn Sci 16, 606–617 (2012).

58. Sadaghiani, S. et al. Alpha-band phase synchrony is related to activity in the fronto-parietal adaptive control network. Journal of Neuroscience 32, 14305– 14310 (2012).

59. Sadaghiani, S. et al. Lesions to the Fronto-Parietal Network Impact Alpha-Band Phase Synchrony and Cognitive Control. Cerebral Cortex 29, 4143–4153 (2019).

60. Klimesch, W., Schimke, H. & Pfurtscheller, G. Alpha frequency, cognitive load and memory performance. Brain Topogr 5, 241–251 (1993).

61. Klimesch, W. EEG alpha and theta oscillations reflect cognitive and memory performance: a rKlimesch, W. (1999). EEG alpha and theta oscillations reflect cognitive and memory performance: a review and analysis. Brain Research Reviews, 29(2-3), 169–195. doi:10.1016/S016. Brain Res Rev 29, 169–195 (1999).

62. Doppelmayr, M., Klimesch, W., Hödlmoser, K., Sauseng, P. & Gruber, W. Intelligence related upper alpha desynchronization in a semantic memory task. Brain Res Bull 66, 171–177 (2005).

63. Klimesch, W., Sauseng, P. & Hanslmayr, S. EEG alpha oscillations: The inhibition-timing hypothesis. Brain Res Rev 53, 63–88 (2007).

64. Gurd, J. M. et al. Posterior parietal cortex is implicated in continuous switching between verbal fluency tasks: An fMRI study with clinical implications. Brain 125, 1024–1038 (2002).

65. Piai, V., Klaus, J. & Rossetto, E. The lexical nature of alpha-beta oscillations in context-driven word production. J Neurolinguistics 55, 100905 (2020).

66. Zakharov, I., Tabueva, A., Adamovich, T., Kovas, Y. & Malykh, S. Alpha Band Resting-State EEG Connectivity Is Associated With Non-verbal Intelligence. Front Hum Neurosci 14, 1–10 (2020).

67. Klepp, A., Niccolai, V., Buccino, G., Schnitzler, A. & Biermann-Ruben, K. Language-motor interference reflected in MEG beta oscillations. Neuroimage 109, 438–448 (2015).

68. Terporten, R., Schoffelen, J. M., Dai, B., Hagoort, P. & Kösem, A. The Relation between Alpha/Beta Oscillations and the Encoding of Sentence induced Contextual Information. Sci Rep 9, 1–12 (2019).

69. Tanaka-Koshiyama, K. et al. Abnormal Spontaneous Gamma Power Is Associated With Verbal Learning and Memory Dysfunction in Schizophrenia. Front Psychiatry 11, 1–9 (2020).

70. Hsieh, L. T., Ekstrom, A. D. & Ranganath, C. Neural oscillations associated with item and temporal order maintenance in working memory. Journal of Neuroscience 31, 10803–10810 (2011).

71. Marshuetz, C. Order information in working memory: An integrative review of evidence from brain and behavior. Psychol Bull 131, 323–339 (2005).

72. Hoffman, K. L. et al. Saccades during visual exploration align hippocampal 3–8 Hz rhythms in human and non-human primates. Front Syst Neurosci 7, (2013).

73. Van Opstal, F., Fias, W., Peigneux, P. & Verguts, T. The neural representation of extensively trained ordered sequences. Neuroimage 47, 367–375 (2009).

74. Polyn, S. M., Norman, K. A. & Kahana, M. J. A Context Maintenance and Retrieval Model of Organizational Processes in Free Recall. Psychol Rev 116, 129–156 (2009).

