## Supplementary material for "Intrinsic Neural Oscillations Predict Verbal Learning Performance and Encoding Strategy Use"

Statistical Analysis and Mediation analysis: To investigate the hypothesis that neural correlates are linked with learning strategies and learning performance, we employed a mediation model incorporating brain oscillatory ROIs as mediators. This model examined both single and serial dual mediator scenarios. Our approach began with a basic linear regression analysis to explore the relationships between various behavioral measures (including semantic clustering, serial clustering, subjective clustering, and learning performance scores) and individual brain ROIs across different frequencies. We selected only those ROIs demonstrating significant associations ($p<0.05$) as potential mediators in our subsequent analysis. The mediation models were then constructed and analyzed using the Hayes PROCESS toolbox. Distinct models were created for subjective and semantic clustering strategies. We applied the bootstrap method (n=5000) for testing both single mediator models and dual serial mediator models, focusing on identifying significant indirect pathways, defined as those showing a non-null range between the lower and upper confidence limits.

**Brain behavior mediates encoding strategies in verbal learning performance**

Resting state theta oscillation mediate subjective clustering to verbal learning performance.

We found that resting theta oscillations enabled us to identify brain regions at rest that are associated, on the one hand, with subjective encoding strategies and, on the other hand, with verbal performance. These models established a causal link between subjective encoding strategies and verbal performance, mediated by specific brain regions. We identified three different patterns with subjective clustering involving only theta oscillations (Figure 7).

The first pattern (Fig 7a) showed a positive regression between subjective clustering and the left sensorimotor regions and the right insula ($r = -.204$; $p = 0.0007$). These latter regions were associated with the left premotor and parietal regions ($r = 1.163$; $p < 0.001$), which, in turn, were negatively associated with verbal learning performance ($r = -4.81$; $p = 0.077$). After introducing the two mediators, the full model (effect = 4.02, $p < 0.001$) comprised a direct path which remained significant (effect = 3.85, $p = 0.0003$), and the indirect path, which included the two mediators (effect = 1.14; bootSE=0.77; boot interval [0.06; 3.08]) (Table 3).

The second pattern (Fig 7b) showed a positive regression between subjective clustering and the left sensorimotor regions and the right insula (r = -.204; p = 0.0007). These latter regions were associated with bilateral occipital and lower limbic temporal regions (r = -0.83; p = 0.0001), which, in turn, were negatively associated with verbal learning performance (r = -6.64; p = 0.0018). After introducing the two mediators, the full model (effect = 4.02, p < 0.001) comprised a direct path which remained significant (effect = 3.97, p = 0.0001), and the indirect path, which included the two mediators (effect = 1.13; bootSE=0.64; boot interval [0.27; 2.73]) (Table 3).

The third pattern (Fig 7c) showed a negative regression between subjective clustering and the left anterior insula, left inferior and anterior temporal and right prefrontal regions (r = -0.21; p = 0.001). These latter regions were associated with bilateral occipital and lower limbic temporal regions (r = 0.42; p = 0.03), which, in turn, were negatively associated with verbal learning performance (r = -5.23; p = 0.004). After introducing the two mediators, the full model (effect = 4.02, p < 0.001) comprised a direct path which remains significant (effect = 3.62, p = 0.003), and the indirect path, which included the two mediators (effect = .48, bootSE=0.36; boot interval [0.01; 1.39]) (Table 3).

<u>*Supplementary figures and table*</u>

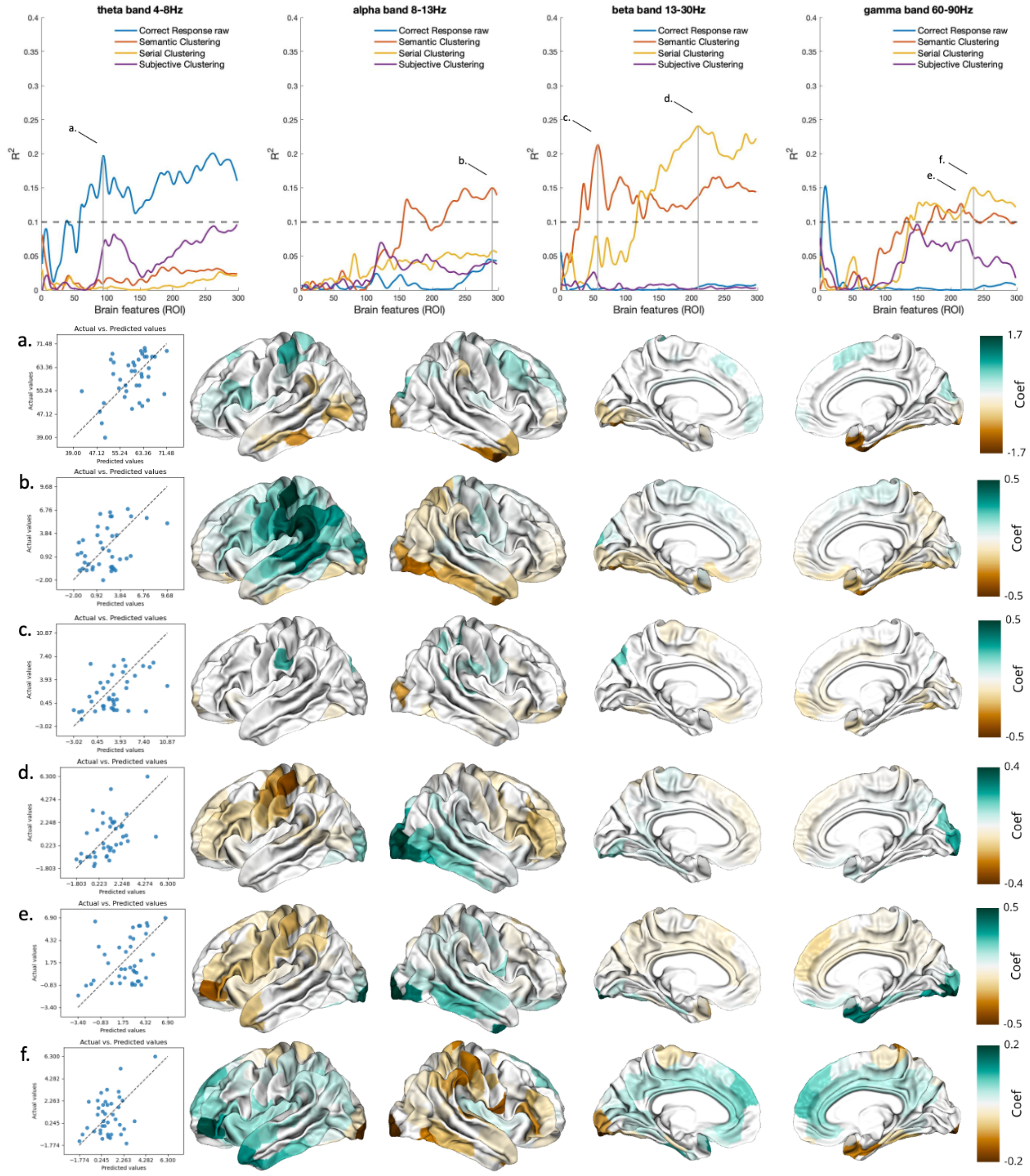


*SF1: The upper panel shows the R² results for the multi-featured prediction model obtained for each brain feature through sequential feature selection (N_ROI; ranging from 2 up to 300 ROIs, i.e., brain space). The lower panel presents a scatter plot illustrating the regression between actual and predicted values based on the best multi-featured prediction model obtained (left side), along with the coefficients extracted relevant to the model for each brain frequency and associated behavioral measure, in*

*relation to data covariance (right side). a) Theta band and Correct Responses; N_ROI = 93; MSE = 54.6; R² = 0.20;*

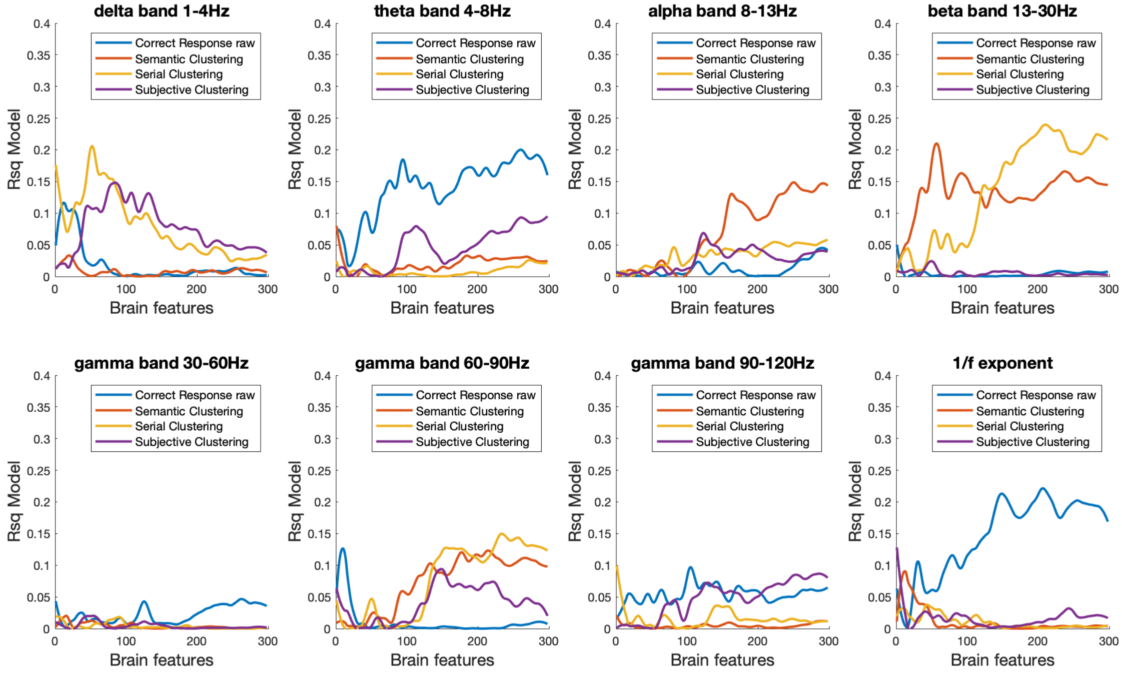


*SF2: Explained variance between actual value and predicted value for each sequential feature selection (ROI), for each frequency band and aperiodic component.*

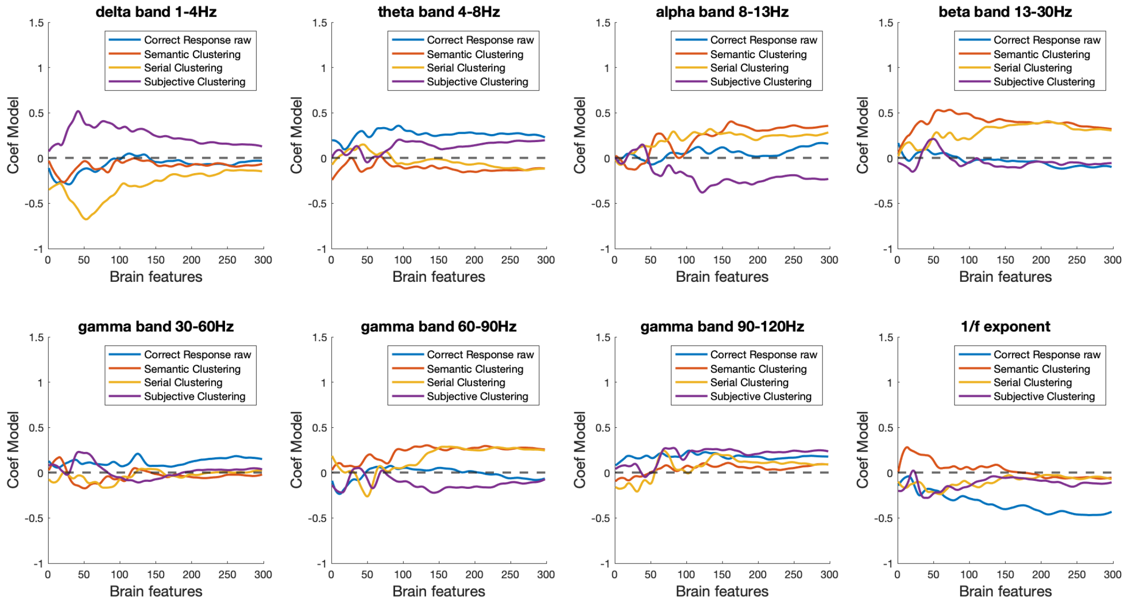


*SF3: Coefficient between actual value and predicted value for each sequential feature selection (ROI), for each frequency band and aperiodic component.*

Table 2: mediation model with single mediator

| Predictors | Subjective Clustering | Semantic Clustering |
|---|---|---|
| Mediators | Gamma band (90-120hz) | Gamma band (90-120hz) |
| Direct | 2.9* | .97* |
| Indirect | 1.05* | .37* |
| Total | 4.0* | 1.43* |

* p<0.05.

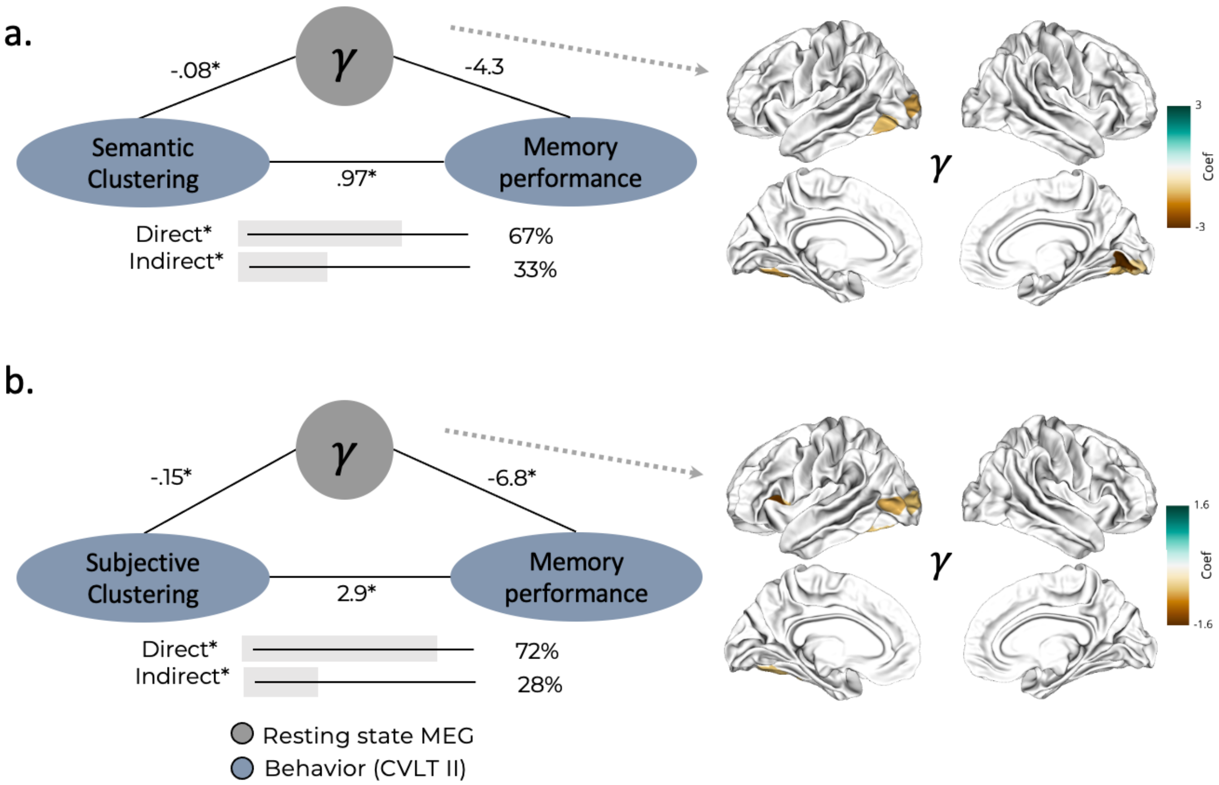


*SF4: Mediation model with single mediators. a) b)*

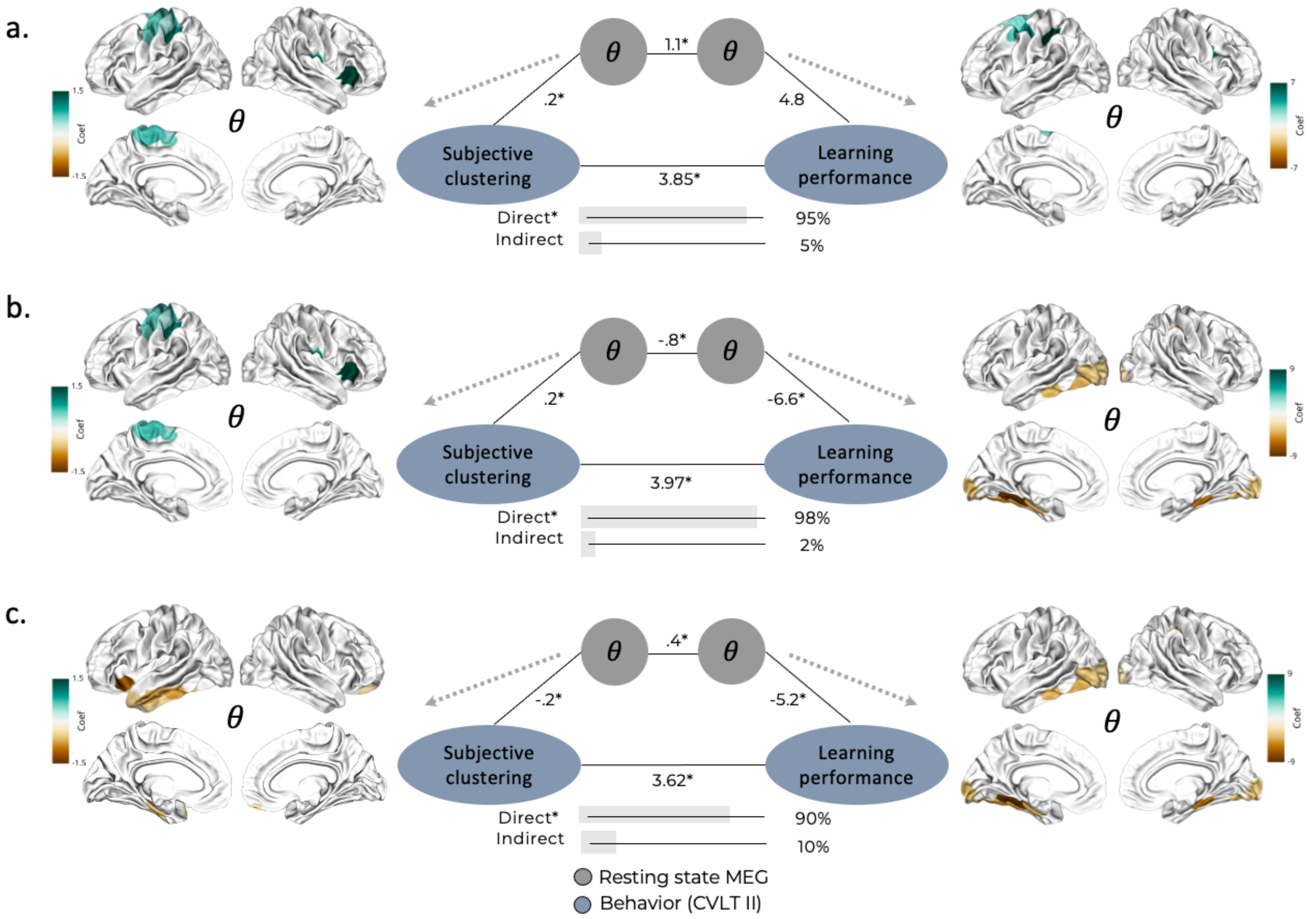


*SF5: Three distinct serial mediation models were employed to predict verbal learning performance (correct responses) using* ***subjective clustering encoding strategies****. On both sides, brain map ROIs were used as mediators. In the middle, the mediation model is depicted with respective pathway coefficients (*: $p < 0.05$).*

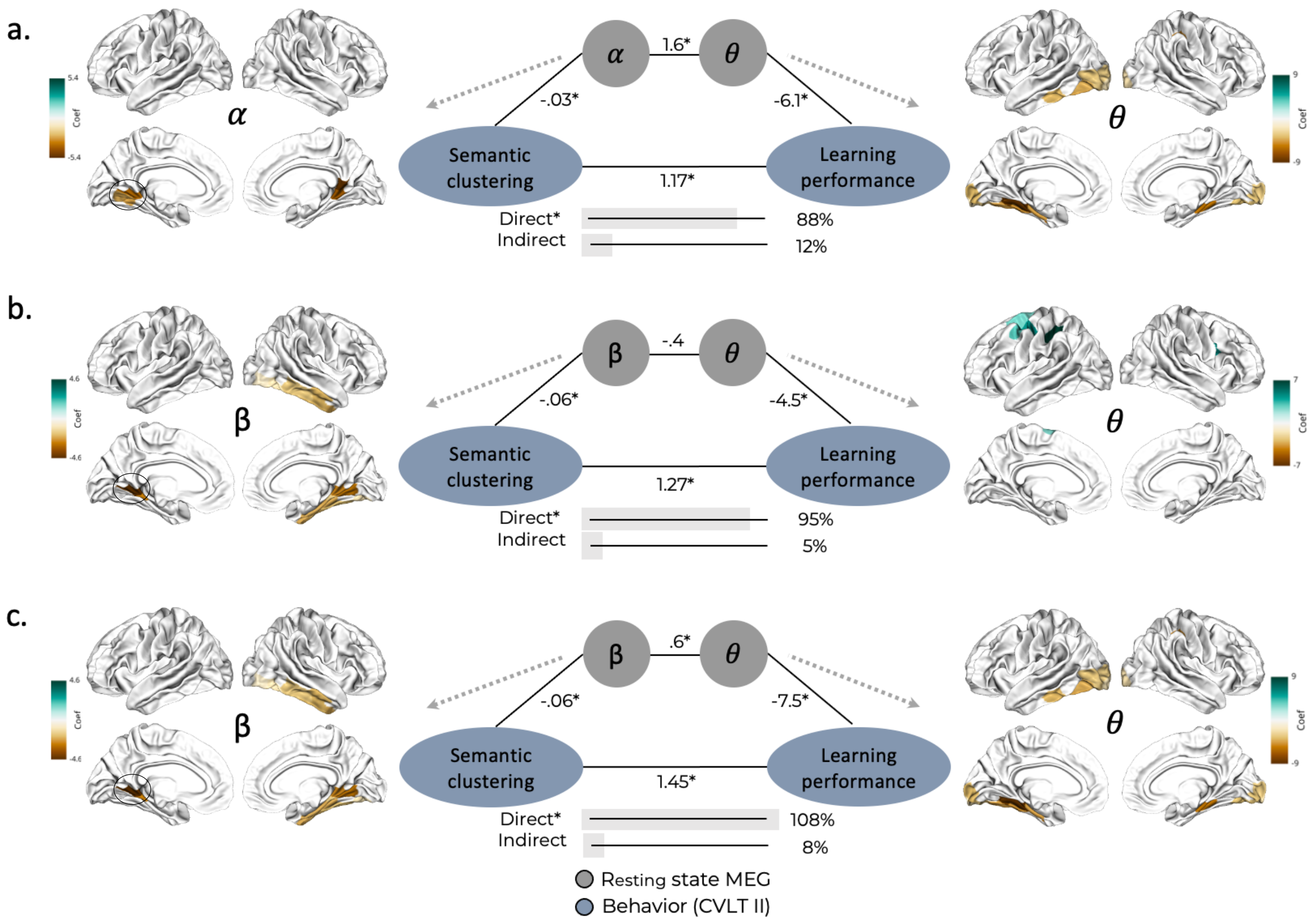


*SF6: Three distinct serial mediation models were employed to predict verbal learning performance (correct responses) using* ***semantic clustering encoding strategies****. On both sides, brain map ROIs were used as mediators. In the middle, the mediation model is depicted with respective pathway coefficients (*: $p < 0.05$).*

<u>Table 3: mediation model results with two mediators</u>

| Predictors | Subjective clustering | | | Subjective clustering | | |
|---|---|---|---|---|---|---|
| Mediators | M1: Theta +<br>M2: Theta + | M1: Theta +<br>M2: Theta - | M1: Theta -<br>M2: Theta - | M1: Alpha -<br>M2: Theta - | M1: Beta -<br>M2: Theta + | M1: Beta -<br>M2: Theta - |
| Direct | 3.85* | 3.97* | 3.62* | 1.17* | 1.27* | 1.45* |
| Indirect | .17 | .04 | .40 | .16 | .06 | -.11 |
| Ind 1 | -.85 | -.84 | -.03 | -.03 | -.19 | -.36 |
| Ind 2 | -.12 | -.25 | -.11 | -.11 | .12 | -.06 |
| Ind 3 | 1.14* | 1.13* | .32 | .32* | .14* | .31* |
| Total | 4.02* | 4.02* | 4.02* | 1.34* | 1.34* | 1.34* |

*p<.05; Ind1: X—Mediator 1—Y; Ind2: X—Mediator 2—Y; Ind3: X—Mediator 1—Mediator 2—Y.

Table 4: Spatial averaging coefficients for each brain behavior pattern

| Network | Beta_Semantic | Beta_Serial | Gamma_Semantic | Gamma_Serial |
|---|---|---|---|---|
| Control | 0.1334 | -0.045 | -0.0487 | 0.0356 |
| Somato-Motor | 0.1189 | 0.0354 | -0.1071 | 0.1457 |
| Dorsal At | 0.1033 | -0.0088 | -0.0672 | 0.0494 |
| Salience | 0.0896 | -0.0353 | -0.076 | 0.0859 |
| Default | 0.0478 | -0.0191 | -0.0276 | 0.0286 |
| Limbic | -0.0782 | 0.0375 | 0.1435 | -0.1497 |
| Visual | -0.2322 | -0.0082 | 0.0873 | -0.1 |
